# A Lung Tumor-on-a-Chip Model Recapitulates the Effect of Hypoxia on Radiotherapy Response and FDG-PET Imaging

**DOI:** 10.1101/2025.07.23.666453

**Authors:** Rohollah Nasiri, Myra Kurosu Jalil, Veronica Ibanez Gaspar, Andrea Sofia Flores Perez, Hieu Thi Minh Nguyen, Syamantak Khan, Sindy KY Tang, Yunzhi Peter Yang, Guillem Pratx

## Abstract

Most solid tumors contain regions of hypoxia that pose a significant challenge to the efficacy of radiation therapy. This study introduces a novel 3D lung tumor-on-a-chip (ToC) model designed to replicate the hypoxic tumor microenvironment *in vitro* while also providing a platform for clinically relevant interventions such as radiotherapy and positron emission tomography (PET) imaging. To simulate the heterogeneous oxygen distribution found in tumors, the ToC model incorporates an oxygen gradient achieved through a straightforward chemical oxygen scavenging system. A unique innovation of this device is the integration of a thin scintillator plate, which enables high-resolution radioluminescence microscopy imaging of tumor metabolism under hypoxia and normoxia conditions using clinically approved PET tracers such as fluorodeoxyglucose (FDG). The response of this hypoxic model to radiation therapy (10 Gy X-ray) demonstrated ∼4-fold higher radioresistance compared to the normoxic ToC model, as assessed by colony formation potential. Additionally, DNA damage observed in the normoxic ToC model was ∼5-fold higher than that in the hypoxic model. Furthermore, the metabolic consumption of glucose was found to mirror the localization of hypoxia, validating the use of this biomarker for planning radiation therapy. The integration of high-resolution radionuclide imaging within ToC models enables on-chip PET imaging and facilitates oncology research and discovery, offering innovative capabilities for the preclinical testing of novel cancer therapies in a clinically relevant environment.

## 1. Introduction

Despite significant progress in treatment, cancer remains one of the leading causes of death worldwide [1]. Half of all patients with cancer undergo radiation therapy (RT) as part of their treatment regimen, which involves targeting tumors with high-energy X-rays while minimizing damage to the surrounding healthy tissues [1][2]. RT is a key treatment for lung cancer, but the heterogeneous nature of the tumor microenvironment (TME) can adversely affect therapeutic outcomes [3][4][5]. A critical aspect of the TME is hypoxia, a condition characterized by low oxygen levels arising due to the abnormal and inefficient vascular networks within tumors [6]. Hypoxic regions within tumors are significantly associated with resistance to radiation and other forms of cancer therapy [7][8][9]. Understanding and targeting hypoxia within the TME is therefore essential for improving the efficacy of cancer treatments.

Current preclinical models, such as static 2D cell culture systems, have little relevance to *in vivo* conditions due to their inability to replicate dynamic and heterogeneous processes, their limited control over the cellular microenvironment, and the absence of critical cell-cell and cell-extracellular matrix (ECM) interactions [10][11]. Similarly, while animal models offer physiological complexity, they also present challenges such as ethical concerns, interspecies differences, low throughput, and high costs, all of which can limit their scalability and accessibility for certain types of studies [11][12][13] . In contrast, 3D tumors-on-a-chip (ToC) are *in vitro* models engineered to replicate essential aspects of the TME, providing physiological relevance and reducing dependence on animal models [15][16][17][18]. These advanced microphysiological systems utilize microfluidics technology to grow tumor cells in 3D within a controlled hydrogel-based microenvironment. Over the past decade, various human tissue constructs have been engineered as models using this technology for studying human physiology and pathology [3][11][20].

Due to these advantages, ToC models have found use in radiotherapy research to study normal tissue and tumor response for different types of radiation sources. Using this technology, the effect of radiation on tumors has been explored using head-and-neck cancer-on-a-chip [21], melanoma cancer-on-a-chip [22], soft tissue carcinoma spheroids-on-a-chip [23], and pancreatic tumor-on-a-chip models [24]. Moreover, the detrimental effects of radiation on normal tissues have been characterized using similar models, including bone-marrow-on-a-chip [25], microvasculature-on-a-chip [26], lung-on-a-chip [27], gut-on-a-chip [28], and also multi-organs on-a-chip (e.g. heart, liver, and bone marrow) [29], all of which show the potential of organ-on-a-chip models in modeling complex responses to RT.

To better understand how the TME affects radiation responses, we developed a 3D lung tumor-on-a-chip in which the oxygen distribution can be easily controlled to induce either hypoxia, normoxia, or a spatial gradient of oxygen across the cell-laden hydrogel. **Figure 1** illustrates the generation of a hypoxia gradient through a perfusion system and the creation of hypoxic conditions using an oxygen scavenger. The chip components are visually represented using different colors to distinguish the normoxia and hypoxia media flows through the side channels, while the central area highlights the tumor tissue culture. The perfusion system, which is connected to the tumor chips, is also illustrated, emphasizing the dynamic environment maintained during the study.

Unique to this study is the integration of a CdWO4 scintillator within the microfluidics device to enable on-chip radioluminescence microscopy (RLM) imaging using clinically relevant radionuclides [28][29][30] . Cancer cells in most tumors rely heavily on glycolysis, a phenomenon that forms the basis for ^18^F-fluorodeoxyglucose positron emission tomography (FDG-PET), a cancer imaging technique used for cancer diagnosis and treatment monitoring. However, the limited spatial resolution of PET (4–8 mm) restricts its ability to resolve the structural and functional heterogeneity of the TME [32]. RLM enables cellular-level metabolic imaging by converting radioactive decays into visible light flashes detectable within a highly sensitive microscope. This approach can be applied to study the complex and heterogeneous TME, evaluate therapeutic responses, and accelerate drug testing in a controlled and clinically relevant platform, integrating the strengths of traditional *in vivo* imaging techniques with advanced tumor-on-a-chip systems.

**Figure 1.**
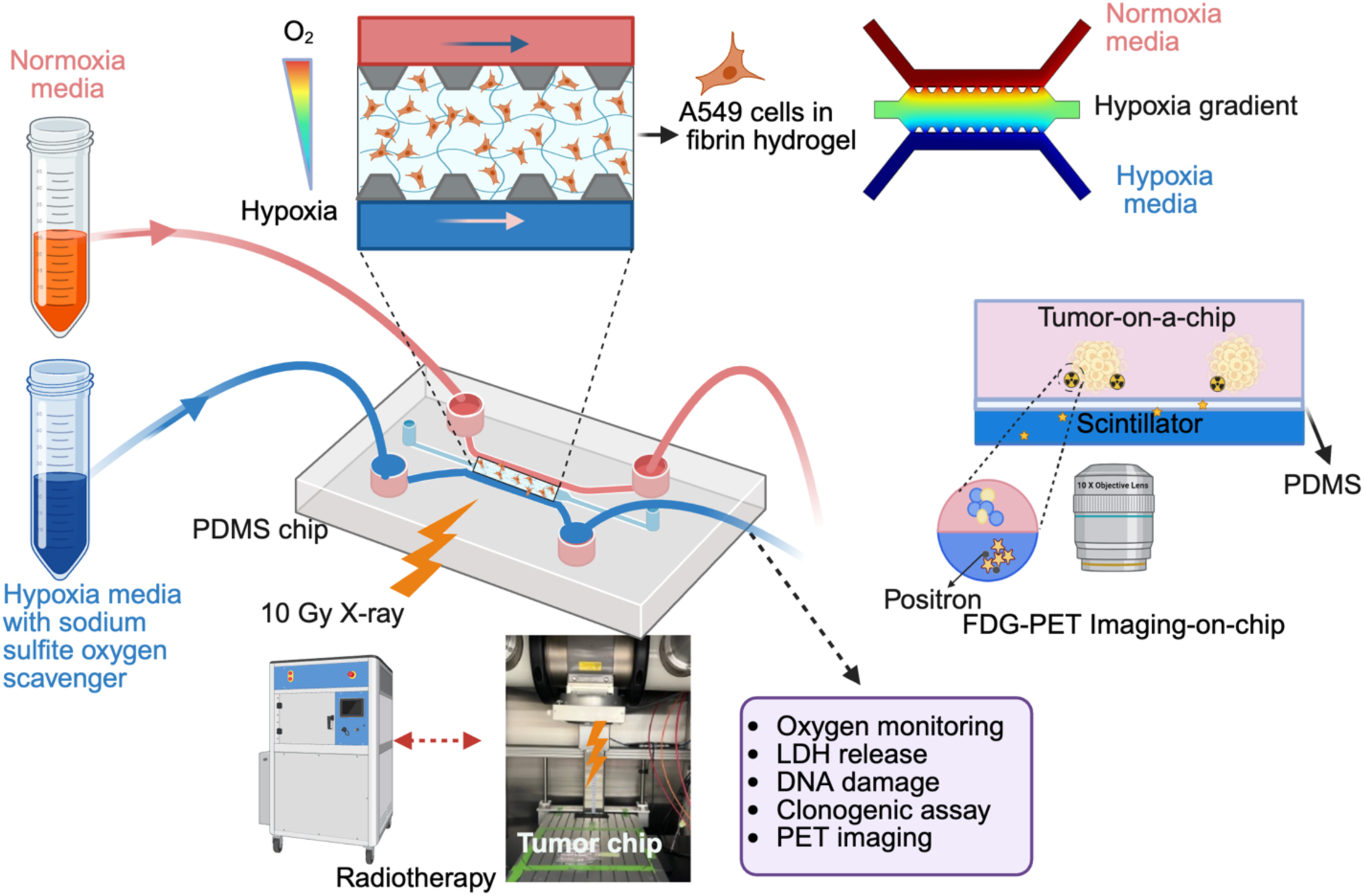
Radiation therapy response of a hypoxic lung tumor-on-a-chip model. Overview of the study, including hypoxia induction, oxygen gradient settings, tumor chip design, radiation delivery, and on-chip radioluminescence and PET imaging using ^18^F-fluorodeoxyglucose. Some elements in this figure were created using BioRender.com.

## 1. Materials and Methods

### Microfluidic chip fabrication

The microfluidic chip was designed with SolidWorks and incorporates a central chamber for housing 3D tumor models, which are separated from side channels by trapezoidal micro-pillars. These side channels are designed to perfuse the tumor models with fresh media, facilitating nutrient delivery and waste removal. We fabricated the master mold from SU-8 2050 photoresist (Kayaku Advanced Materials) on a silicon wafer (University Wafer) using standard photolithography. To achieve a nominal feature height of 250 µm while minimizing height variation, we performed two spin coats at a final spin speed of 1450 RPM, each to produce a 125 µm layer of photoresist. We soft-baked the wafer for 5 minutes at 65 °C and 12 minutes at 95 °C after the first spin coating step, and 5 minutes at 65 °C and 25 minutes at 95 °C after the second spin coating step. We used a direct write machine (Durham Magneto Optics MicroWriter ML3) to expose the features of the mold at 390 mJ/cm2 with 5 µm resolution. After exposure, we baked the wafer for 5 minutes at 65 °C and 20 minutes at 95 °C, developed the master mold, and performed a hard bake for 5 minutes at 200 °C. The measured feature height was 250 µm ± ∼10% (223.6 µm - 277.8 µm). We passivated the master mold using trichloro(1H,1H,2H,2H-perfluorooctyl)silane (catalog no. 448931, Sigma Aldrich) prior to use.

The microfluidic channels were fabricated by soft lithography. Polydimethylsiloxane (PDMS) (Sylgard 184 silicone elastomer base and curing agent) was prepared by mixing the PDMS prepolymer with a curing agent at a ratio of 10:1 w/w. After thorough mixing to avoid air bubbles, the PDMS mixture was poured onto the patterned silicon wafer, degassed under vacuum, and cured at 80°C for 2 hours. Once cured, the PDMS replica was peeled off the silicon mold. Inlets and outlets for media perfusion were created by punching 1.2 mm diameter holes through the PDMS using a biopsy punch (Unicore Punch Kit, Sigma). The PDMS chip was subsequently bonded to a glass slide to create a sealed microfluidic device using a corona discharger (Electro-Technic Products) to activate the surfaces for 2 minutes, ensuring a strong bond between PDMS and glass.

### On-chip radioluminescence microscopy

For ^18^F-fluorodeoxyglucose (FDG) imaging using RLM, the microfluidic device incorporates a scintillator at its bottom in place of the traditional glass slide. Hence, an additional preparation step involves spin-coating a thin PDMS layer onto a CdWO4 scintillator (2 cm × 2 cm × 0.5 mm; Kinheng Crystal Material (Shanghai) Co., Ltd.) to facilitates attachment of the microfluidics device to the radiation-sensing scintillator [33]. A solution of PDMS diluted in tert-butyl alcohol (1:4 w/w) is spin-coated at 2200 RPM for 30 minutes to create a uniform layer with a thickness below 5 µm. After coating, the PDMS layer is cured at 80°C for two hours to ensure strong adhesion and stability. The coated scintillator is then bonded to the PDMS channel using plasma treatment. Both surfaces are exposed to plasma for 2 minutes using a corona discharge system before bonding. This plasma treatment modifies the surface chemistry by converting Si–CH₃ groups to Si–OH, which facilitates the formation of covalent Si–O–Si bonds upon contact. This chemical bonding enhances the adhesion between the PDMS chip and the PDMS-coated scintillator, optimizing the setup for efficient RLM imaging.

#### Cells

This study was performed using A549 cells, a human alveolar basal epithelial adenocarcinoma cell line. Purchased from ATCC, these cells were cultured in F12-K medium supplemented with 10% fetal bovine serum (FBS) and 1% penicillin-streptomycin. Cultures were maintained in a humidified atmosphere with 5% CO2 at 37°C and passaged at 70-80% confluency to ensure optimal growth conditions.

### Tumor-on-a-chip model fabrication

Prior to cell culture, the chips were treated with plasma to enhance hydrophilicity. They were then immersed in ethanol for two hours, and the channels were flushed three times with ethanol. Subsequently, the chips were rinsed three times with phosphate-buffered saline (PBS) to remove the ethanol. A 3D ToC model was developed by incorporating A549 lung cancer cells into a fibrin hydrogel scaffold. To do so, a fibrinogen solution was prepared at 10 mg/mL in PBS, incubated at 37°C for 30 minutes until complete dissolution, and sterilized through a 0.2 µm filter. To create the hydrogel, the fibrinogen solution was diluted with cell culture media to achieve a final fibrin concentration of 3 mg/mL and a cell density of 3 million cells/mL. The mixture was gently mixed to ensure uniform cell distribution. Thrombin was added at a final concentration of 1 U/mL to initiate polymerization for ∼2 minutes before seeding the mixture into the chip. The chips were then incubated for 10 minutes to complete gelation, after which media was added to the side channels. The tumor model was perfused with media at a flow rate of 3 µL/min using a peristaltic pump (BT100F-1 Intelligent Dispensing Peristaltic Pump with Pump Head DG6-24), mimicking dynamic conditions.

### Hypoxia induction

Hypoxia was induced by supplementing the media with sodium sulfite, an oxygen scavenger that lowers dissolved oxygen levels, and cobalt nitrate, which stabilizes hypoxia-inducible factor-1 alpha (HIF-1α) to mimic hypoxia signaling. A hypoxia gradient was created by perfusing one of the side channels with hypoxic media and the other with normoxic media. Additionally, normoxic and hypoxic conditions were induced by perfusing both inlets with either normoxic or hypoxic media, respectively. The hypoxic medium was prepared by dissolving sodium sulfite (Na₂SO₃; Sigma-Aldrich-7757-83-7) and cobalt (II) nitrate hexahydrate (Thermo Scientific Chemicals) in cell culture media. Sodium sulfite (100 mg) was dissolved in 10 mL DMEM and 2.93 mg cobalt (II) nitrate was prepared as a 2 mM stock solution. The final hypoxic media contained 1% sodium sulfite (w/v) and 100 µM cobalt nitrate.

### Hypoxia imaging

Hypoxia was monitored using the Image-iT Green Hypoxia Reagent (Thermo Fisher). A 2 µM concentration of the reagent was added to hypoxic and normoxic media, and cells were incubated under hypoxic conditions for 4 hours. After incubation, cells and chips were washed with PBS to remove any excess dye. Imaging was performed using a Leica microscope (DMi8) with GFP filter settings to visualize hypoxic cells. This method enables qualitative and spatially resolved detection of cellular hypoxia in live-cell cultures.

### Radiation treatment

Radiation treatment of the ToC model was performed using an X-ray irradiator (Precision X-ray XRAD 320). Tumor chips were placed inside the X-ray machine and exposed to a dose of 10 Gy. The X-ray machine was operated at 320 kV and 12.5 mA and equipped with a treatment filter (1.5 mm Al, 0.25 mm Cu, and 0.75 mm Sn). The machine was previously calibrated according to standard TG61 methodology [34].

### Immunofluorescence (IF) staining for DNA damage visualization

To visualize DNA damage post-radiation, cells were incubated for 45 minutes to allow DNA repair to take place. DNA double-strand break damage was assessed by staining for γ-H2AX, a well-established molecular marker of DNA damage [38][39][40]. Cells were fixed with 4% paraformaldehyde (PFA, ThermoFisher) for 15 minutes at room temperature, followed by permeabilization with 0.1% Triton X-100 (ThermoFisher) and blocking with 3% bovine serum albumin (BSA, Fisher Scientific) in PBS. They were then incubated overnight at 4°C with a primary antibody against γ-H2AX (anti-phospho-histone H2A.X (Ser139), Sigma-Aldrich, Catalog # 05-636-I) at a 1:100 dilution in 1% BSA staining buffer. The next day, cells were washed and incubated for 1 hour at room temperature with an Alexa Fluor 488-conjugated secondary antibody (donkey anti-mouse IgG (H+L), Alexa Fluor™ 488, ThermoFisher, Catalog # A-21202) at a 1:100 dilution. Following three washes with PBS, nuclei were stained with Hoechst 33342 (5 µg/ml) for 20 minutes, then washed again. The samples were imaged using fluorescence microscopy to visualize DNA damage foci. Fluorescent intensity was analyzed using ImageJ to quantify DNA damage for each sample group.

### F-actin staining

Cells were fixed, permeabilized, and blocked as described previously. Hoechst was used to stain cell nuclei. For F-actin staining, Alexa Fluor™ 555 Phalloidin (ThermoFisher) was applied at a 1:200 dilution (v/v) in a staining buffer containing 1% BSA. The cells were incubated with the staining solution at room temperature for staining of F-actin filaments.

### Cell death assay

Dead cells were identified using SYTOX Green (Invitrogen; NucGreen Dead 488 ReadyProbes Reagent). One drop of the reagent was added per 1 ml of media and applied to the cells. The dye penetrates cells with compromised membranes, binding to nucleic acids and emitting green fluorescence, while live cells with intact membranes exclude the dye, remaining unstained. This method enables sensitive and accurate detection of dead cells inside the chip.

### Alamar Blue viability assay

To evaluate the viability of cancer cells under different hypoxia levels, an Alamar Blue assay was conducted in a 24-well plate. A549 cells were seeded onto at a cell density of 200×10^3^ cells per well and grown for 24 hours. Cells were exposed to varying oxygen scavenger concentrations for 1 day, and a 10% Alamar Blue solution (ThermoFisher) in cell culture media was added to each well. The plate was incubated for 3 hours at 37°C. After incubation, absorbance was measured at 570 nm using a plate reader to assess cell viability. This approach allows for the quantification of cell viability, providing insight into how different hypoxic conditions induced by oxygen scavenging affect cell viability and cytotoxicity.

### Clonogenic assay

After exposing the ToC model to 10 Gy of X-ray radiation under normoxic, hypoxic, or gradient conditions, the cells were extracted from the chips using 0.25% trypsin to digest the fibrin gel. The hydrogel was incubated with trypsin at 37°C for 10–15 minutes with gentle agitation to ensure uniform digestion. Following enzymatic digestion, trypsin was neutralized with serum-containing medium, and the cell suspension was centrifuged at 300 g for 5 minutes. The pellet was then washed once with PBS or fresh medium before proceeding with downstream applications. Subsequently, the cells were counted and seeded into 6-well plates at a density of 200 cells/well for non-irradiated samples and 2000 cells/well for irradiated samples to assess their colony-forming ability. After a 14-day incubation period, the cells were fixed with 4% paraformaldehyde and stained with 0.1% crystal violet. Colonies consisting of more than 50 cells were visualized and counted, providing quantitative data on cell survival and clonogenic potential. Cells from unirradiated ToCs served as reference to estimate the surviving fractions of cells receiving RT under different oxygen levels.

### Lactate dehydrogenase (LDH) release

LDH release, indicative of cell membrane damage and cytotoxicity, was measured 24 hours after X-ray exposure. Cell culture supernatants were collected and transferred to a 96-well plate for analysis. An LDH detection kit (CytoTox 96, Cytotoxicity assay, Promega) was used according to the manufacturer’s instructions. Absorbance was measured using a plate reader at 490 nm wavelength. Increased absorbance values correspond to higher levels of LDH release, indicating greater cell membrane damage due to X-ray exposure.

### Expression of hypoxia-related genes

To assess the expression of hypoxia-related genes (CA-IX and GLUT3), quantitative polymerase chain reaction (qPCR) analysis was performed. The process began with cell lysis using an appropriate lysis buffer (BioRad Total RNA mini kit) to extract total RNA. A total of 800-1000 ng of starting RNA was used as a template for cDNA synthesis, using the iScript kit from BioRad. qPCR was set up with PowerUp SYBR Green Master Mix, according to manufacturer’s instructions. In brief, each sample contained 5 µl of SYBR Green Master Mix (2x), 1 ng of cDNA and 800 nM of forward and reverse primers (each). Nuclease free water was added to a total reaction volume of 10 µl. The primers were CA-IX, Glut 3, and GAPDH as an internal control gene (see Table 1 for the forward and reverse primer sequences). Non-template controls for each gene were also prepared simultaneously. Samples were run in duplicates. The qPCR was run in a QuantStudio 6 using a fast-cycling mode consisting of UDG activation (50°C, 2 min), activation (95°C, 2 min), and 40 cycles of denaturation (95°C, 1 s) and anneal/extend (60°C, 30 s). This was followed by a melt curve stage. Results were analyzed using the delta delta CT method.

**Table 1.**
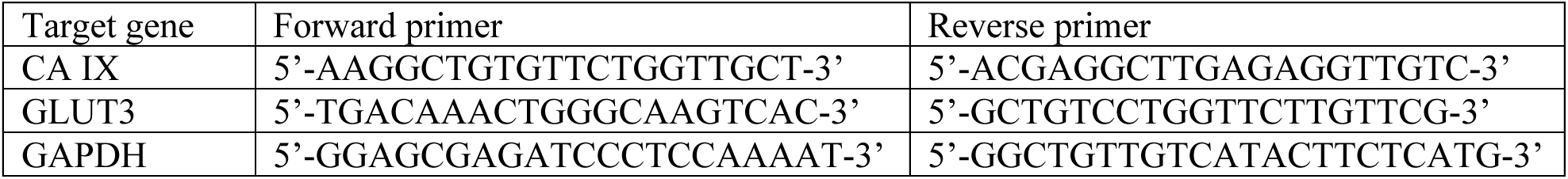
Primer sequences used for qPCR analysis of target genes (CA IX and GLUT3) and the housekeeping gene (GAPDH).

### Numerical simulations

Numerical simulations of fluid flow and oxygen concentration in the ToC model were performed using a Finite Element Method (FEM) solver (COMSOL Multiphysics 4.4.a). The process involves modeling laminar flow and the Navier-Stokes equations for fluid dynamics, alongside concentration equations for oxygen transport. The chip model includes two inlets with equal flow rates but differing oxygen concentrations (0 and 0.2 mol/m^3^ O2), simulating physiological conditions and the formation of an oxygen gradient across the device chamber. The model included both free-flow regions (side channels) and a porous medium (central hydrogel chamber), with fluid flow governed by the Navier-Stokes equations in the side channels and Darcy’s Law in the hydrogel region to account for permeability effects. The simulation parameters for the porous hydrogel were set assuming a permeability of 10⁻^12^ m² and porosity of 0.9 to reflect average gel properties for fibrin [35][36]. Boundary conditions at the outlet assumed atmospheric pressure (P=0 Pa; relative pressure), ensuring realistic fluid dynamics behavior. No-slip conditions are applied to the chip walls, assuming zero velocity at the solid boundaries. This simulation setup enables detailed analysis of oxygen distribution and flow patterns within the microfluidic chip, crucial for optimizing experimental conditions.

### Bulk FDG uptake measurements

Bulk FDG uptake in cells under normoxic and hypoxic conditions was measured using an automated gamma counter (PerkinElmer Wizard, Revvity). FDG was synthesized at the Stanford radiochemistry facility using an on-site cyclotron by standard nucleophilic fluorination of mannose triflate. Cells in ToC models were incubated with perfusion of FDG (1 mCi/mL) in glucose-free media (DMEM, 1% FBS, 0.1% P/S) (with a flow rate of 3 μL/min) for 60 minutes to allow uptake via glucose transporters and trapping via hexokinase, reflecting glucose metabolism as measured clinically with PET/CT. After incubation, the ToC models were washed three times with PBS. Cells were then extracted, and viable cell numbers were counted using trypan blue staining (Gibco, Catalog #15250061) and a cell counter (Countess 3 Automated Cell Counter, ThermoFisher). The cells were subsequently lysed to release intracellular FDG and metabolites. The lysates were transferred to flow cytometry glass tubes for gamma counting. The gamma counter quantified radioactive emissions as counts per minute (CPM), providing a direct and clinically relevant measure of FDG uptake and cellular metabolic activity. FDG uptake values were normalized to the corresponding cell number.

### On-chip radioluminescence microscopy

High-resolution radioluminescence microscopy of FDG uptake was performed in ToC models cultured within special PDMS microfluidic chips that incorporated a 0.5 mm-thick CdWO4 scintillator at the bottom of the device. After 4 days of culture, the ToC models were incubated with FDG (1 mCi/mL) in glucose-free DMEM medium for 60 minutes to facilitate uptake and metabolism of the tracer, reflecting glucose metabolism on a spatial level. Post-incubation, the chips were washed three times with PBS to remove residual FDG. They were then mounted onto a standard glass coverslip (0.1 mm thick, Fisher Scientific) and placed on the RLM stage for imaging. Scintillation events originating from single radioactive decays were recorded during imaging by acquiring a sequence of 2,000 luminescence frames, each acquired with a 500 ms exposure time. In addition, standard brightfield images were acquired to focus the microscope, select a suitable field of view, and provide co-registered structural images of the ToC. Frames capturing radioactive decay events were processed using a custom MATLAB scripts to reconstruct and analyze FDG uptake, enabling precise visualization of metabolic activity [31]. This innovative approach offers a robust platform for characterizing the impact of microenvironmental conditions and therapeutic interventions on tumor metabolism according to a clinically relevant endpoint, the uptake of FDG.

### PET-CT imaging of ToC models

To compare RLM against standard PET imaging, the uptake of FDG by ToC models was measured using a PET/CT scanner (GNEXT, Sofie Biosciences). PET images were acquired with a 10-minute acquisition time and the standard energy window (350 to 650 keV). CT images were captured using a 2-minute scan with 80-kVp beam energy. PET data were reconstructed using the OSEM method provided by the scanner’s vendor, and images were analyzed with OsiriX software. Maximum intensity projection was used to display PET data, optimizing slice thickness to match that of the chip model.

### Statistical analysis

All group comparisons in this study were performed using Prism software. One-way ANOVA was used when analyzing a single independent variable, and two-way ANOVA was used when two independent variables were involved. Tukey’s post-hoc test was applied for multiple comparisons. Statistical significance was defined as follows: *P < 0.05, **P < 0.01, ***P < 0.001, and ****P < 0.0001.

## 3. Results

First, we demonstrate the ability of the oxygen scavenger to induce robust and durable hypoxia *in vitro*. A549 cells were cultured under hypoxic and normoxic conditions for 24 hours, then treated with Image-iT Green Hypoxia Reagent. As shown in **Figure 2A**, cells under hypoxic conditions displayed strong green fluorescence signal, confirming the successful establishment of hypoxia. In contrast, no signal was observed in cells cultured under normoxic conditions. In addition, qPCR analysis revealed significant upregulation of two key hypoxia-related genes, CA IX and GLUT3, under hypoxic conditions compared to normoxia (**Figure 2B**). CA IX expression increased approximately 4-fold, while GLUT3 showed an ∼8-fold increase under hypoxia relative to normoxia. To evaluate the potential cytotoxicity of the oxygen scavenger, a cell viability assay and an Alamar Blue assay were performed to identify an acceptable concentration range (**Fig. S1** and **Fig. S2**). Both assays showed a decreasing trend in cell viability with increasing oxygen scavenger concentrations. While these effects could be due to the scavenger itself, they are more likely a consequence of hypoxic conditions. In light of these findings, we selected a 1% sodium sulfite concentration, as it effectively induces hypoxia while maintaining acceptable cell viability. This concentration also aligns with previous studies utilizing sodium sulfite for hypoxia creation [6][37]. Collectively, these findings confirm the effectiveness of the oxygen scavenger in creating hypoxic conditions.

**Figure 2.**
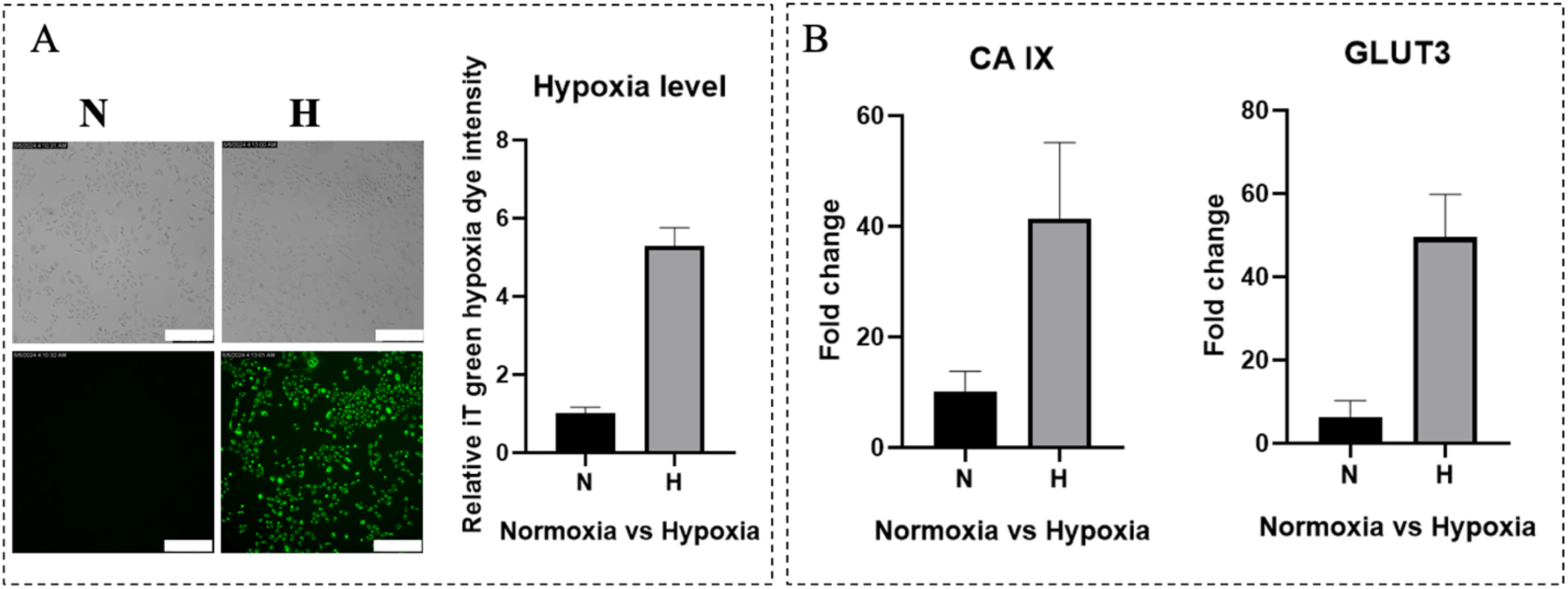
Induction and monitoring of hypoxia in A549 cells. (A) Hypoxia (H) (with 1% sodium sulfite) and normoxia (N) conditions monitored using Hypoxia Green dye. (scale bar = 300 µm). (B) qPCR analysis of key hypoxia-related genes CA IX and GLUT3. Data represent mean ± SD from n = 3 independent samples.

### Tumor-on-a-chip model development

We then investigate the use of the oxygen scavenger in a ToC model. **Figure 3** shows the chip design and fabrication process. The chip was designed using SolidWorks (**Fig. 3A-i**), then the master mold was fabricated via photolithography (**Fig. 3A-ii**), and the final device along with its inlets and outlets for media perfusion was molded using soft PDMS (**Fig. 3A-iii**). The 3D lung tumor model, cultured within the central channel of the microfluidic chip, is depicted in **Fig. 3B-**This central region is flanked by two side channels and is separated from them by trapezoidal microposts, which effectively maintain surface tension and prevent media or gel spillage into adjacent channels during loading. The design eliminates the need for co-flow, enabling precise localization of the tumor model within the central compartment. Additionally, fluorescence imaging of F-actin and Hoechst staining was conducted to provide clear visualization of the tumor cells within the microfluidic chip (**Fig. 3B-ii)**. **Fig. 3B-iii** illustrates the use of stainless-steel plugs to seal the inlets and outlets of the central chamber after seeding the tumor cells. **Fig. 3B-iv** shows the ToC devices connected to the perfusion system for continuous media flow. **Fig. 3B-v** uses food coloring to depict the three different conditions being modeled in this study: normoxic, hypoxic, and hypoxia.

**Figure 3.**
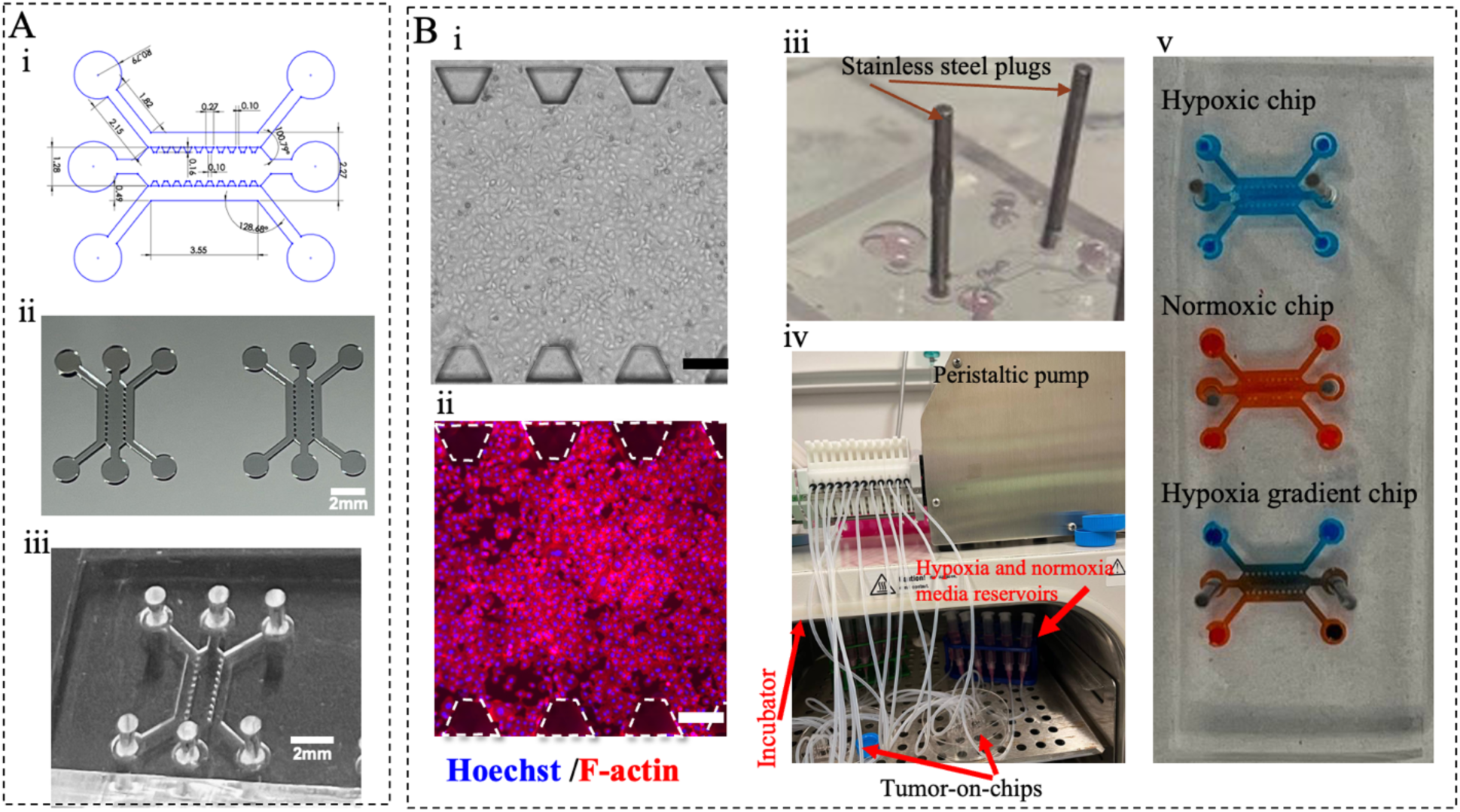
Tumor-on-a-chip development. (A). (i) Chip design (dimensions are in mm), (ii) Fabricated master mold, and (iii) final PDMS chip. (B). (i) Brightfield image of the 3D ToC model, (ii) Hoechst/F-actin staining (scale bar=300 µm), (iii) Sealing the tumor chip with stainless steel plugs before perfusion, (iv) Perfusion system setup for the ToC model. (v). Representative images of chips under three different conditions: normoxia, hypoxia, and oxygen gradient.

### Numerical simulations for flow dynamics and oxygen concentration

Numerical simulations of fluid flow and oxygen concentration distribution within the ToC model were performed using COMSOL Multiphysics at flow rates of 1, 3, and 6 µL/min. **Figure 4A-i-ii** illustrates the fluid velocity distribution across the chip, while **Figure 4A-iv** presents velocity profiles across the central channel (along the mid-line depicted in **Fig. 4A-iii**) for different flow rates. A similar velocity profiles in the side channels arise from the symmetric design of the chip, while minimal velocity magnitude is observed in the middle channel. All channels exhibit a parabolic velocity profile from wall to wall, with the zero-velocity observed at the channel walls corresponding to the no-slip boundary condition. The oxygen concentration distribution was modeled using the diffusion equation, with the diffusivity constant of oxygen set to D=1.5×10^−9^ m^2^/s. In this simulation, one inlet was assigned an oxygen concentration of 0 mol/m³ (hypoxia), while the other was set to 0.2 mol/m³ (mM) (normoxia), reflecting physiological conditions. **Figure 4B i-ii** depicts the resulting oxygen concentration gradient within the chip, successfully demonstrating the transition from normoxia to hypoxia across the central chamber. The corresponding oxygen concentration profile along the central width of the channel is also shown, emphasizing the gradient formation as a function of the flow rate in the lateral channels (**Figure 4B iii**). Shear stress distribution across the chip was also evaluated for 3 µL/min flow rate (**Figure 4C-i**), showing higher values along the walls and minimal shear stress in the central region of the channel where cells are cultured. The simulations also captured the volumetric average shear stress distribution in the central channel of the chip (tumor region), as shown in **Fig. 4C-ii**, for three different flow rates. The results demonstrate an increasing trend in average shear stress with increasing flow rate. With the physical properties of water considered—density of 1000 kg/m³ and viscosity of 0.001 Pa·s—these numerical simulations provide valuable insights into fluid dynamics, oxygen distribution, and shear stress behavior within the ToC model. It is important to note that these simulations do not include cellular oxygen consumption. Therefore, the oxygen gradients shown here represent the effect of physical diffusion and flow-based transport, independent of metabolic activity.

**Figure 4.**
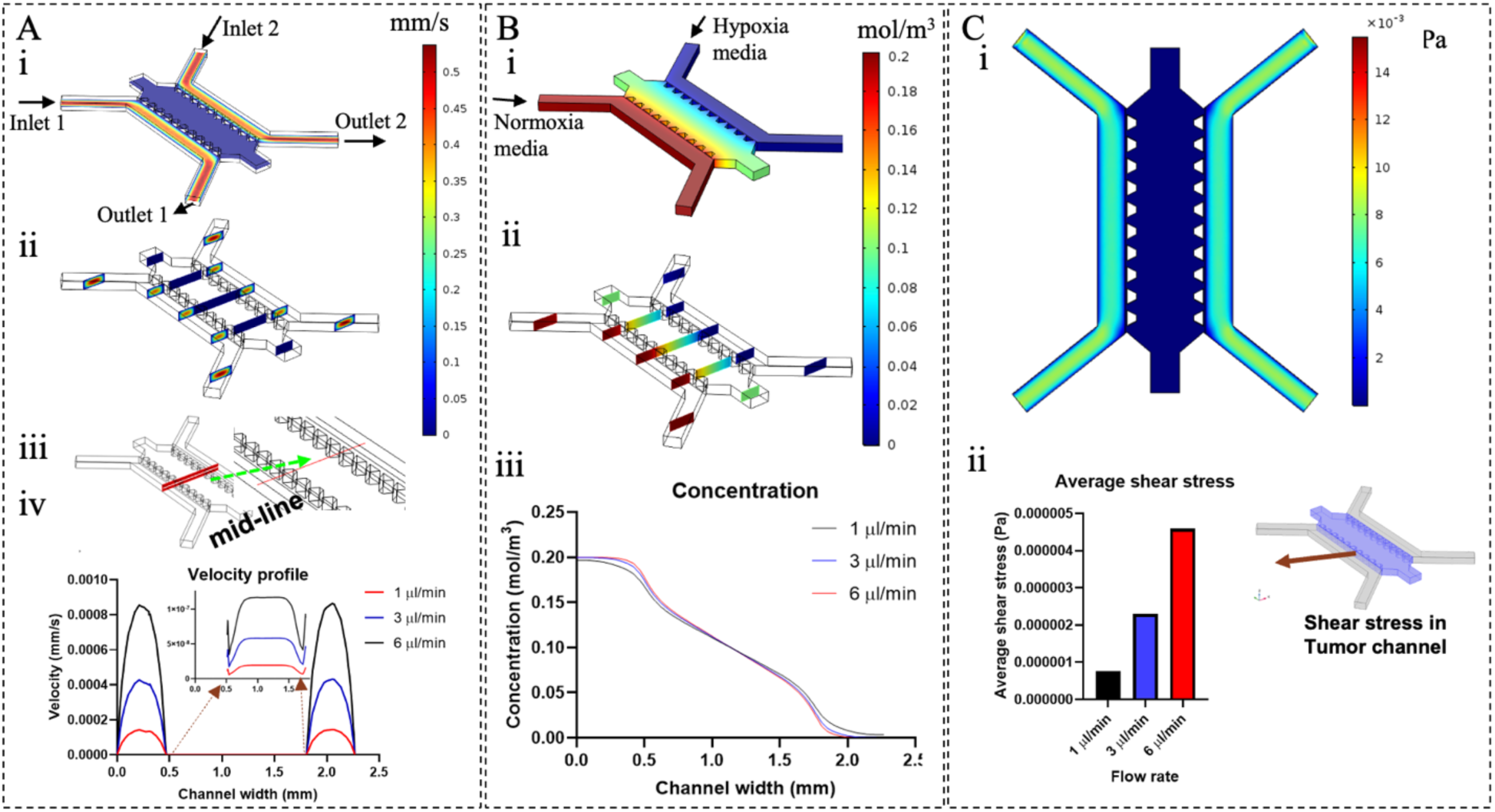
Numerical Simulation for Tumor-on-a-Chip. (A). i. Velocity distribution along the horizontal mid-plane of the chip shows higher velocity magnitudes in the side channels compared to the middle channel (flow rate of 3 µL/min). ii. Velocity distribution across different vertical cut planes highlights the highest velocity in the center of each channel cross-section due to the parabolic velocity profile typical of laminar flow in microchannels (at 3 µL/min). iii. Definition of the mid-line. iv. Velocity profile distributions along mid-line, spanning the side channels and middle channel. (B). i. Oxygen concentration distribution within the channel and mixing pattern at a flow rate of 3 µL/min. Concentration distribution across different cut planes along the channel length. iii. Concentration profiles along the mid-line of the central channel, showing a gradient in oxygen concentration for three flow rates (1, 3, and 6 µL/min). (C). i. Shear stress distribution across the mid-plane of the channel at a flow rate of 3 µL/min. ii. Average shear stress values at different flow rates within the central tumor-on-a-chip model.

### Hypoxia monitoring in ToC model

The induction of hypoxia in the ToC model using an oxygen scavenger was assessed using a fluorogenic hypoxia sensor (Image-iT Green Hypoxia Reagent). **Figure 5A** displays the tumor chip models under three conditions: normoxia, hypoxia, and hypoxia gradient. The fluorescence signal from the hypoxia sensor reflects oxygen levels under each condition. The normoxic chip exhibits minimal fluorescence, confirming the absence of hypoxia. In contrast, the gradient chip shows a gradual increase in fluorescence intensity across the channel, reflecting the established oxygen gradient. The fully hypoxic chip demonstrates maximum fluorescence, confirming widespread hypoxia. These findings were verified by simulating the device with COMSOL, assuming the same experimental conditions (e.g. flow rate of 3 µL/min; **Figure 5B**). To characterize the hypoxia gradient, the fluorescence of the hypoxia sensor was quantified using ImageJ within six regions of interest (ROIs; shown in Fig. 5A-i). **Figure 5C** displays relative hypoxia levels in these ROIs, showing low and spatially uniform fluorescence within the normoxia chip, high fluorescence within the hypoxia chip, and a monotonous increase in fluorescence along the lateral direction in the hypoxia gradient chip. These observations qualitatively align with numerical simulation results shown in **Figure 5B**, validating the successful induction of hypoxia in chip and our ability to modulate oxygen distribution across the central ToC culture chamber. Quantitative alignment between simulation and experiment is not expected, as the fluorogenic hypoxia dye is intended to highlight regions of hypoxia below a set threshold of oxygen (approximately 5%) and is not designed to report quantitative pO₂ values. Hypoxia monitoring in other replicates (**Fig. S3**) further confirms the reproducibility of our ToC model in generating consistent, physiologically relevant hypoxic zones.

**Fig. 5.**
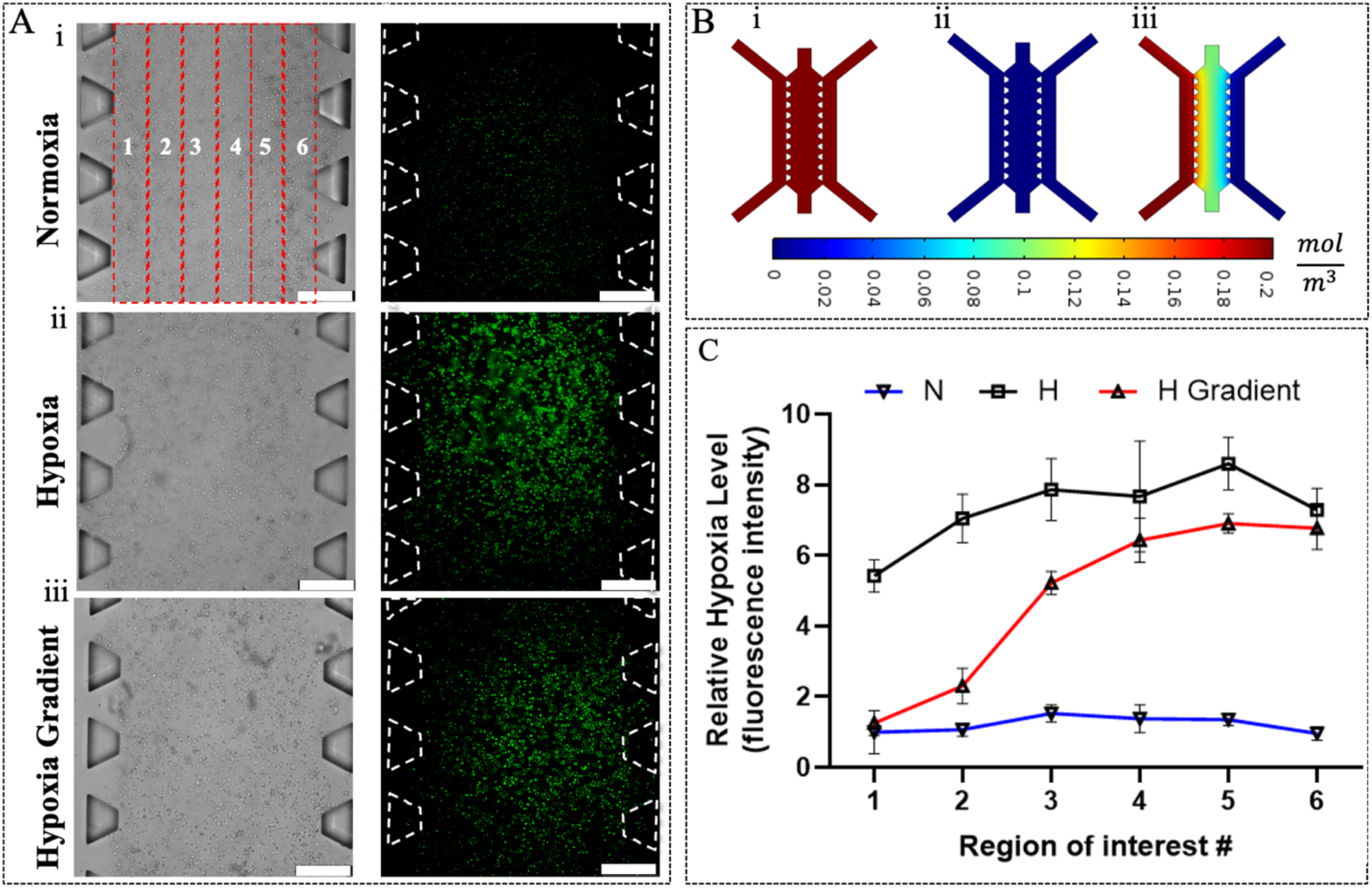
(A) Brightfield and fluorescence images reflecting hypoxic conditions in i. normoxia, ii. hypoxia, and iii. hypoxia-gradient chips. (B) Simulated oxygen distribution for different oxygen environments (i. Hypoxia, ii. normoxia, and iii. oxygen gradient). (C) Quantified relative hypoxia levels in the selected regions of interest (ROIs) (shown as red boxes in A-i) for different oxygen conditions. Scale bar = 300 µm. Data represent mean ± SD from *n* = 3 independent samples.

### Assessment of radiotherapy response in ToC model

After establishing ToC models with controllable hypoxia conditions, radiation therapy was applied using a preclinical X-ray irradiator (10 Gy; **Figure 6A)**. To evaluate the response of the ToC models to radiation therapy, we performed a clonogenic assay, a key technique for quantifying the surviving fraction post-radiation. Unlike other viability assay, the clonogenic assay is not influenced by viable cells that are permanently arrested. **Figure 6B** shows the ability of cells harvested from ToC models to form colonies after radiation therapy, while **Figure 6C** quantifies the survival fraction under different hypoxia conditions based on these data. Cells cultured in fully hypoxic conditions for 24 hours exhibited significant radioresistance, forming a higher number of colonies than cells under normoxic conditions after exposure to radiation. For a radiation dose of 10 Gy, the survival fraction in hypoxia was ∼4 times higher than in normoxia, indicating a significant reduction in radiation-induced cell death under low-oxygen conditions. Furthermore, the hypoxia gradient chip displayed an intermediate level of radioresistance, suggesting that even partial hypoxia in solid tumors can result in significant radioresistance when assessed on a whole-tumor level.

Another endpoint for quantifying the response of ToC models to radiation therapy is the measurement of LDH release into the supernatant or perfused media 24 hours post-radiation. LDH release serves as an indicator of cell cytotoxicity since the enzyme is sequestered inside the cytosol of viable cells but released upon membrane damage. To perform this assessment, the media in the chips were replaced with fresh normoxic media immediately after radiation therapy, and the chips were cultured for an additional 24 hours. Subsequently, the supernatants were collected for LDH analysis using a plate reader. LDH release values were subsequently normalized to the number of cells under each condition.

After radiation therapy, LDH release was 3.1 times lower in the hypoxia chip and 2.2 times lower in the oxygen gradient chip relative to the normoxic chip (**Figure 6D**). This finding aligns with the clonogenic assay results and highlights the protective role of hypoxia in enhancing both cell survival and clonogenic survival following radiation therapy.

**Figure 6.**
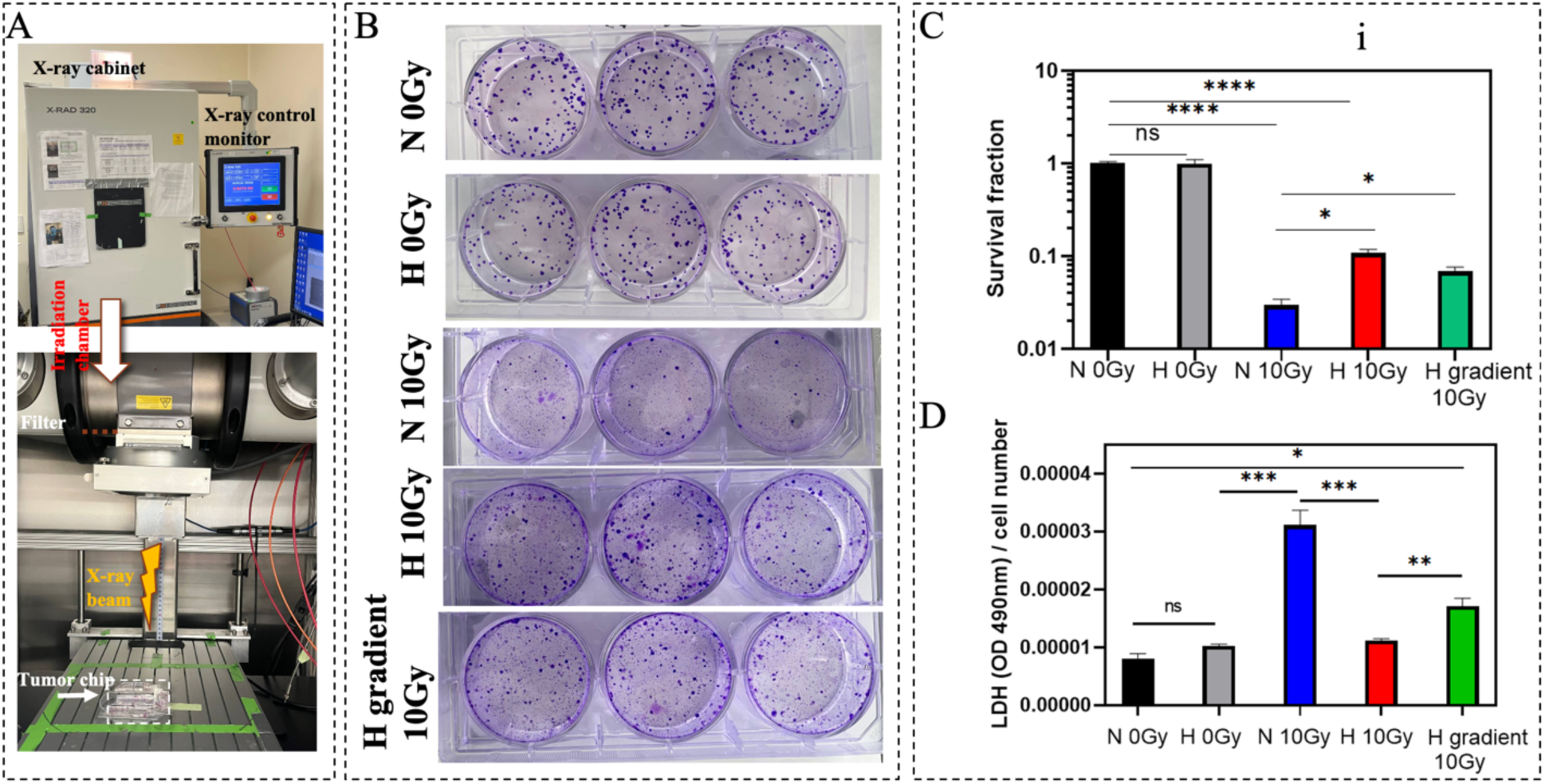
(A) Irradiation of ToC models with a preclinical X-ray irradiator. (B) Clonogenic survival assay depicting colony formation under various conditions: normoxia 0 Gy, hypoxia 0 Gy, normoxia 10 Gy, hypoxia 10 Gy, and hypoxia gradient 10 Gy (from top to bottom). (C) Clonogenic survival rates and (D) LDH release levels under different oxygen conditions following radiation therapy. Data represent mean ± SD from n = 3 independent samples.

DNA damage is critical in mediating the effects of radiation therapy on tumor cells. Damage occurs almost instantaneously upon radiation exposure, while the subsequent DNA repair processes take place over minutes to hours. To assess DNA double-strand break damage in tumor cells, the phosphorylation of histone H2AX (γH2AX) was detected through immunostaining. **Figure 7A** illustrates DNA damage in ToC models measured 45 min after exposure to 10 Gy X-ray radiotherapy, under various conditions, including normoxia, hypoxia, and hypoxia gradient. The intensity of the green fluorescence reflects the extent of DNA damage in different regions of the tumor models within the chip. The highest fluorescent signal was achieved after irradiating the chip with 10 Gy under normoxic conditions, indicating higher DNA damage compared to the hypoxia and hypoxia gradient chips. Conversely, minimal damage is observed in non-irradiated control chips regardless of oxygen conditions. To quantify DNA damage on a spatial level, the tumor region (central channel) was divided into six ROIs (**Figure 7A-i**), and fluorescence intensity was quantified using ImageJ (**Figure 7B**). The results show a uniform pattern of DNA damage for normoxia and hypoxia chips, whereas the hypoxia gradient chip exhibits a gradual increase in damage, reflecting the underlying oxygen gradient. According to these results, DNA damage is increased 5-fold under normoxia compared to hypoxia, highlighting the significant impact of the hypoxic microenvironment on radioresistance. Notably, in the hypoxia gradient ToC model, minimal DNA damage is observed in the hypoxic region, while the normoxic region exhibit the highest damage. These findings further emphasize the oxygen-dependent nature of radiotherapy response, particularly in heterogeneous tumors. **Fig. S4** includes an additional replicate for each condition.

**Fig 7.**
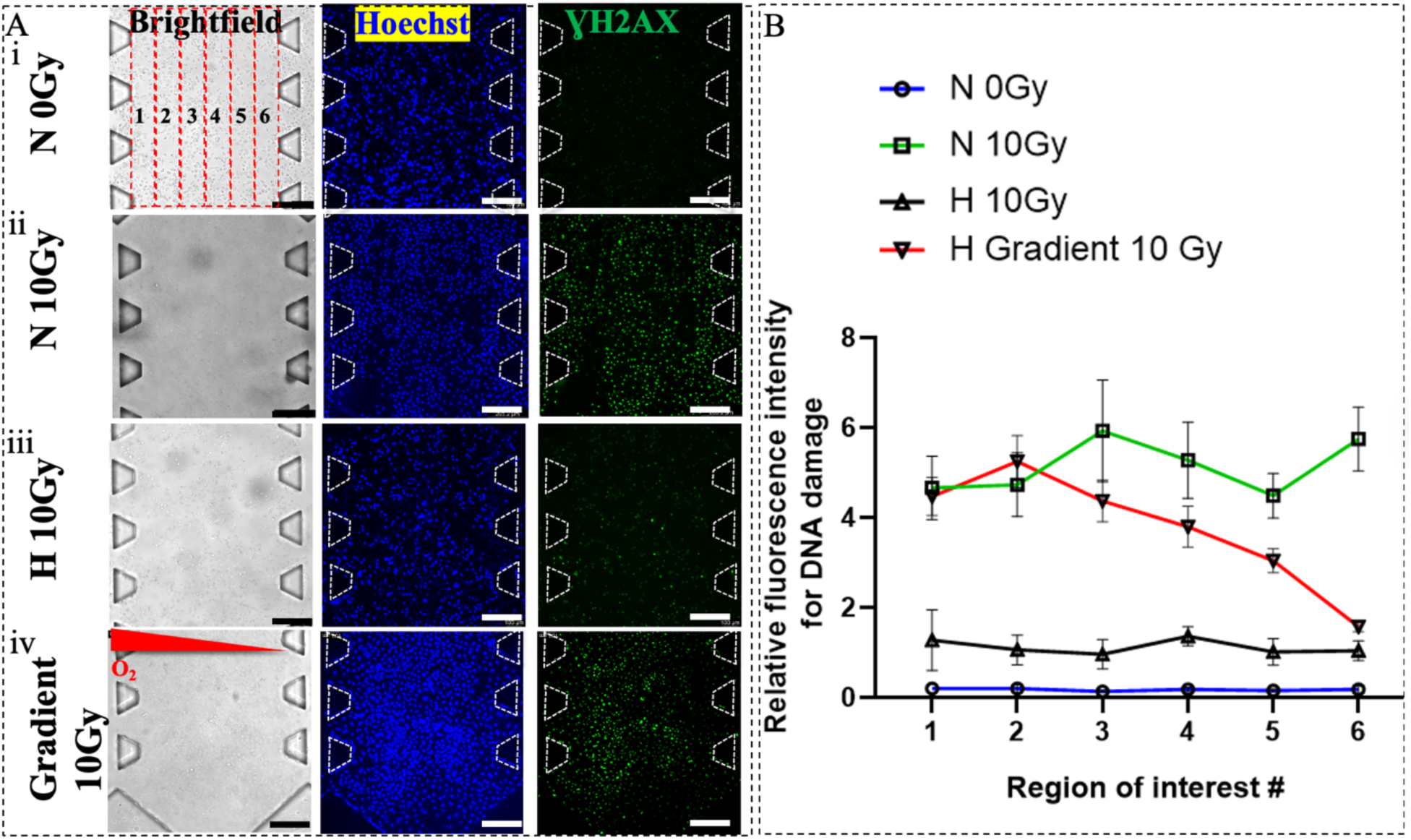
DNA damage detection in the tumor-on-a-chip model. (A) Brightfield, Hoechst, and γ-H2AX staining images for i. Normoxia (0 Gy), ii. Normoxia (10 Gy), iii. Hypoxia (10 Gy), and iv. Hypoxia gradient (10Gy). Scale bar=300 µm. (B) DNA damage quantification across different conditions and ROIs. Data represent mean ± SD from n = 3 independent samples.

### Integration of tumor-on-a-chip model with CdWO4 scintillator for radioluminescence imaging

Modern radiation therapy relies on anatomical and functional imaging to guide the delivery of the radiation treatment and monitor its efficacy. In particular, PET imaging with^18^F-flurodeoxyglucose (FDG) serves as an effective method to map glucose utilization in tumors and identify tumor regions at greater risk for local relapse after radiation. The association between radiotherapy failure and elevated FDG uptake arises from the fact that high FDG uptake is indicative of tumor cells that are either rapidly proliferating or hypoxic, both of which contribute to increased resistance to radiation [41].

A key feature of the ToC device used in this study is the integration of a CdWO4 scintillator (**Figure 8A**) to enable high-resolution radioluminescence imaging of FDG uptake within the 3D culture chamber. This approach provides a platform for high-resolution imaging and metabolic assessment of tumor cells under controlled microenvironmental conditions using FDG, a clinically relevant radiotracer, as the main endpoint. FDG was introduced into the tumor chip to label tumor cells and evaluate their utilization of glucose as a function of oxygen levels. Ionizing positrons emitted following the radioactive decay of ^18^F atoms propagate through the ToC chamber. Upon reaching the scintillator underneath, the positrons trigger the emission of visible light flashes that are captured by a sensitive EMCCD camera coupled to a high-efficiency microscope. **Figure 8B-C** shows the RLM system and EMCCD camera, respectively, that were used in this study to image the ToC model. Once introduced into the tumor chip, FDG is taken up by cells via glucose transporters, then phosphorylated and trapped by hexokinase. As the first step in the glycolysis pathways, hexokinase is upregulated under hypoxia. The accumulation of FDG is then quantified through a reconstruction algorithm and displayed over a conventional brightfield image of the ToC model.

To image glucose metabolism, 1 mCi/ml FDG was perfused through the lung ToC model for 1 hour, and its uptake by tumor cells was monitored using RLM. **Figure 8D** shows RLM imaging results of ToC models under different oxygen conditions: hypoxia, normoxia, and a hypoxia gradient. In these images, higher FDG uptake appears as brighter spots and corresponds to clusters of metabolically active cells. Due to the limitations of RLM in imaging multiple chips simultaneously, imaging was performed sequentially at ∼30-minute intervals, thus the measured uptake values were corrected for radioactive decay (but not for higher efflux at later imaging timepoints). Compared to the normoxia chip, the hypoxia chip exhibited higher FDG uptake. This pattern aligns with the well-established Pasteur effect, where cells under hypoxia shift toward anaerobic glycolysis, leading to increased glucose consumption and elevated FDG uptake in low-oxygen regions. Additionally, the spatial gradient of FDG uptake mirrored the gradient of oxygen, supporting the notion that FDG-PET can highlight areas of the tumors more resistant to radiotherapy that should be targeted with increased dose, a concept known as “dose painting”. Moreover, similar results were obtained in ToC models containing A549 tumor cells cultured in 2D under a gradient of hypoxia (**Fig. S5**).

To independently validate the FDG uptake of cancer cells under normoxic, hypoxic, and hypoxia-gradient conditions, we performed bulk gamma counting using A549 cells cultured in ToC models (**Fig. 8E-i-ii**). After a 1-hour perfusion of 1 mCi/mL FDG, cells were extracted, counted, and their FDG uptake quantified as counts per minute (CPM) via gamma counting. To ensure accurate comparisons, uptake values were normalized to cell number. **Fig. S6** shows FDG uptake values under different conditions before normalization. Since the oxygen scavenger affects cell viability and proliferation, we normalized FDG uptake to cell numbers to enable accurate comparisons across conditions. As shown in **Figure 8E-iii**, the normalized FDG uptake was highest under hypoxia (∼65% higher than normoxia), while the hypoxia-gradient chip also showed elevated uptake (∼30% higher than normoxia). These findings highlight the utility of our ToC platform in evaluating tumor cell FDG uptake under physiologically relevant oxygen conditions. This finding is consistent with the higher uptake of hypoxic tumors and is due to the upregulation of glycolysis under anaerobic conditions.

Finally, as a comparison to RLM, we also utilized a preclinical PET/CT scanner to assess FDG uptake in a tumor-on-a-chip model (in normoxia condition; **Figure 8F**). The CT image shows the basic features of the chip, such as inlets and outlets, whereas the PET image indicates regions of metabolically active tumor cells within the chip. A key advantage of PET/CT is that it eliminates the need for integrating scintillators into the microfluidic device while also enabling high-throughput simultaneous imaging of multiple chips under different treatment conditions. Another advantage is that PET/CT is commercially available compared to RLM. The downside, however, is a ∼20-fold decrease in spatial resolution, which prevents accurate measurement of microscopic-scale heterogeneity within the ToC model.

The integration of radionuclide imaging within the ToC model represents a significant advancement, providing a quantitative and visual tool for assessing tumor metabolism and microenvironmental variations based on a clinically relevant endpoint. This novel approach opens avenues for studying tumor biology and therapy responses with high precision and resolution, while mirroring the clinical workflow.

**Fig. 8.**
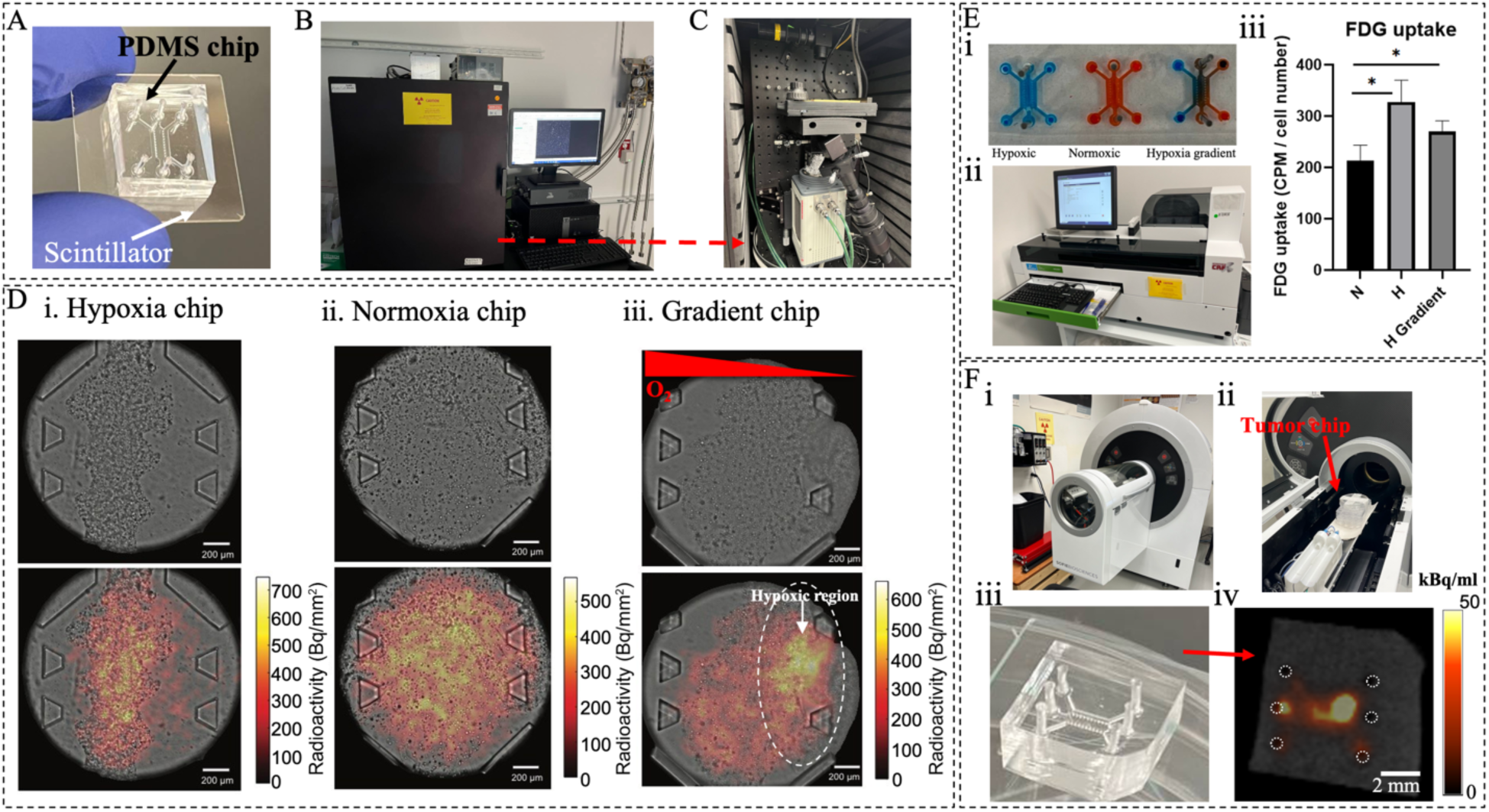
(A). Tumor-on-a-chip integrated with a scintillator for radioluminescence microscopy (RLM) imaging. (B). RLM system enclosed in light-tight box. (C). RLM system, including EMCCD camera, low-light optics, epifluorescence module, and specimen stage. (D). Brightfield images and RLM imaging showing FDG uptake in the tumor-on-a-chip models under i. hypoxia, ii. normoxia, and iii. hypoxia gradient conditions. (E). (i) Illustration showing ToC models under hypoxic, normoxic, and hypoxia gradient conditions for quantification of FDG uptake. (ii) Gamma counter for FDG uptake measurements. (iii) FDG uptake values of ToC models, normalized to cell number, highlighting differences in glucose utilization under different oxygen conditions. Data represent mean ± SD from n = 3 independent samples. (F). i. PET/CT system. ii. Tumor chip mounted in the PET/CT system for imaging. iii. Tumor-on-a-chip. iv. PET/CT imaging showing FDG uptake in tumor chip within the central cell culture chamber .

## Discussion and Conclusion

In this study, we demonstrate a 3D ToC model that incorporates heterogeneous hypoxic conditions, one of the key challenges in treating solid tumors with radiotherapy. Through various assays, we confirmed that even a small region of hypoxia can dramatically increase tumor radioresistance, as shown by increased tumor cell survival and decreased DNA damage and cell death. The observed pattern of DNA damage mirrored the spatial gradient of oxygen under heterogeneous hypoxic conditions, further highlighting the potent role of oxygen in modulating the effect of radiation on cells.

Additionally, this model enabled the monitoring of FDG uptake via RLM imaging, showing the potential of incorporating high-resolution radionuclide imaging in microfluidic 3D cell culture systems. High FDG uptake is associated with increased risk of local relapse post-therapy [41]. In the context of our study, FDG uptake mirrored the incidence of hypoxia in the 3D model, which was itself associated with increased radioresistance. Overall, the integration of radionuclide imaging within this ToC microfluidic platform marks an important advancement in studying tumor responses to therapies using clinically relevant imaging biomarkers such as FDG uptake. RLM is compatible with all PET tracers and could be used as a platform both for the development of new PET tracers, or to assess therapeutic interventions using clinically validated imaging endpoints.

Our study used an oxygen scavenger to induce hypoxia within the microfluidics device [37]. Other available methods include hypoxia chambers, nitrogen infusion, and gas-impermeable microfluidic chips [42][6]. Optimizing media composition, establishing oxygen gradients, and real-time monitoring can further enhance physiological relevance. In our study, sodium sulfite provided a rapid and controllable method to induce hypoxia, offering more flexibility than hypoxia chambers or gas-impermeable chips by enabling localized oxygen depletion without specialized equipment. The establishment of hypoxia gradients with this technique is a valuable *in vitro* model of the diffusion of oxygen that occurs *in vivo* from capillaries into surrounding tumor tissues [43]. While the ToC model used for this study only included a single cell type, these models can also incorporate endothelial cells and fibroblasts to create perfusable vascular networks in the central culture chamber [44][45]. These more advanced systems could be applied to model the role of vascular damage in the response of tumors to radiation or chemotherapy. Furthermore, the incorporation of perfusable vasculature into these models provides a new avenue for studying how the properties of the vascular networks (density, tortuosity, etc) influence the delivery of drugs and imaging agents to the tumor, resulting in different response and uptake. Tumor organoids derived from patient samples can also be incorporated in place of immortalized cancer cell lines to individualize these models according to the unique biological characteristics of each patient’s tumor [46][47][48].

To improve the generalizability of our ToC model, we evaluated the response to radiotherapy in another lung cancer cell line (H1299), under various oxygen conditions. As shown in **Fig. S7**, the model was developed with a similar configuration as previously described, embedding H1299 cells in a 3 mg/mL fibrin hydrogel. Following the establishment of hypoxia and exposure to 10 Gy X-ray radiation, DNA damage was assessed via γH2AX staining and was significantly higher under normoxic conditions than under hypoxia. Additionally, a spatial gradient of DNA damage was observed in the hypoxia-gradient chip, further confirming the oxygen-dependent nature of radiotherapy response and supporting the broader applicability of our platform (**Fig. S8**). FDG uptake was also measured in H1299 cells cultivated in ToC system, as previously described. Hypoxic cells exhibited higher FDG uptake than normoxic cells, with intermediate levels observed in the gradient chip (**Fig. S9)**. Together, these findings validate the versatility of our model across different lung cancer cell types.

Beyond this study, the developed ToC platform can be employed to create various tumor models, enabling precise control and real-time monitoring of key tumor microenvironment parameters under radiotherapy, including pH (via optical sensors), cytokine secretion (via multiplexed assays), oxygen gradients, and nutrient/metabolite dynamics (via biosensors or mass spectrometry). These capabilities, along with compatibility with patient-derived cells, make the platform a versatile tool for modeling complex TME dynamics and improving the translational relevance of preclinical radiotherapy studies.

In conclusion, using a hypoxic lung ToC model, we confirmed the important role that oxygen distribution plays in mediating tumor radioresistance. This model incorporated an inorganic scintillator to enable high-resolution radionuclide imaging uptake via RLM. Overall, the integration of radionuclide imaging within microphysiological models marks an important advancement in studying tumor responses to therapies, paving the way for more precise investigations using clinically relevant imaging endpoints.

## Acknowledgments

Research reported in this publication was supported by the National Cancer Institute of the National Institutes of Health (NIH) under Award Number R01CA268514. The content is solely the responsibility of the authors and does not necessarily represent the official views of the National Institutes of Health. This manuscript is the result of funding in whole or in part by the NIH. It is subject to the NIH Public Access Policy. Through acceptance of this federal funding, NIH has been given a right to make this manuscript publicly available in PubMed Central upon the official date of publication, as defined by NIH.

## Data availability

The data that support the findings of this study are available from the corresponding author upon reasonable request.

