## Supplementary file for "A Lung Tumor-on-a-Chip Model Recapitulates the Effect of Hypoxia on Radiotherapy Response and FDG-PET Imaging"

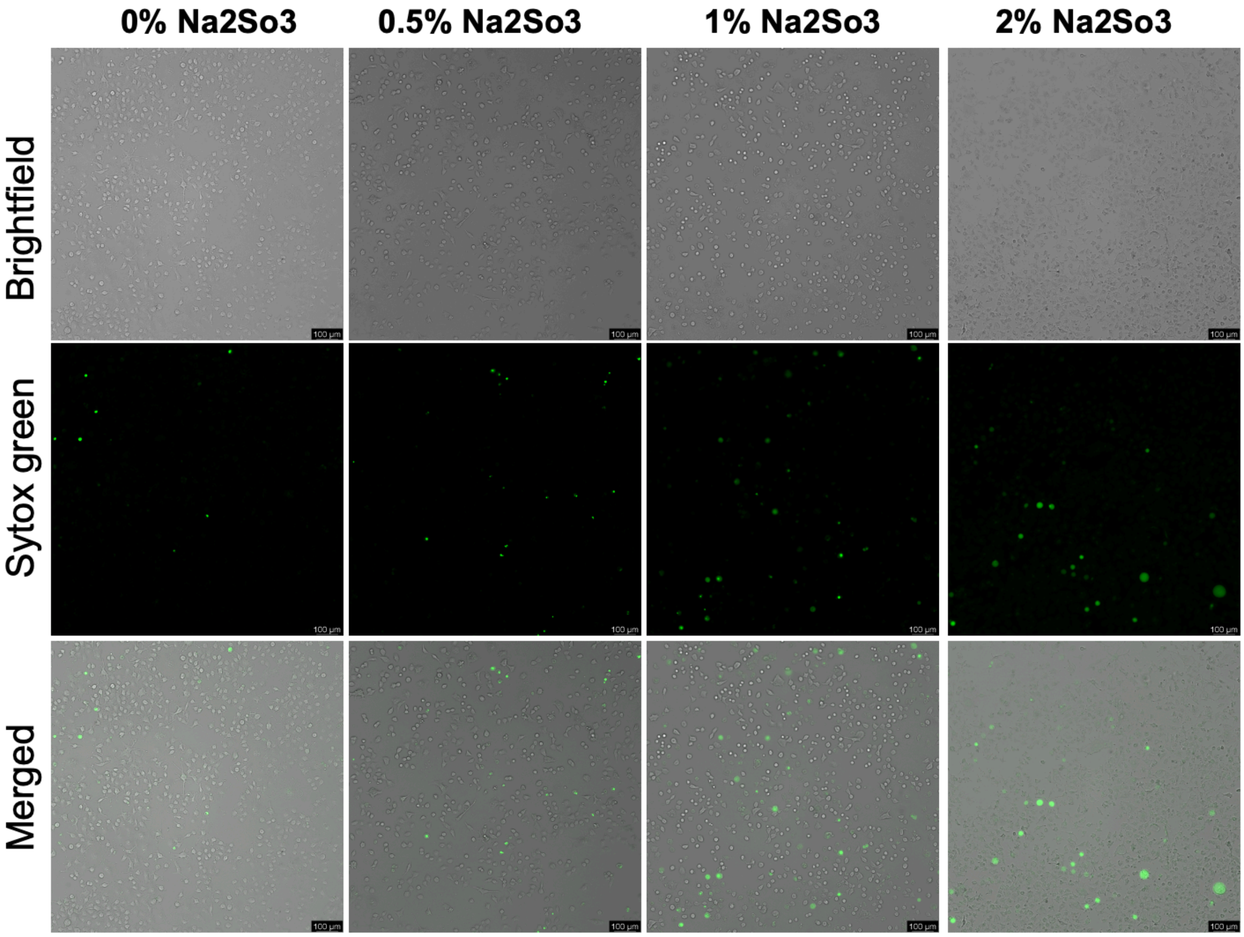


**Fig. S1**. Cell viability at different concentrations of the oxygen scavenger. Green fluorescence indicates dead cells stained with Sytox Green.

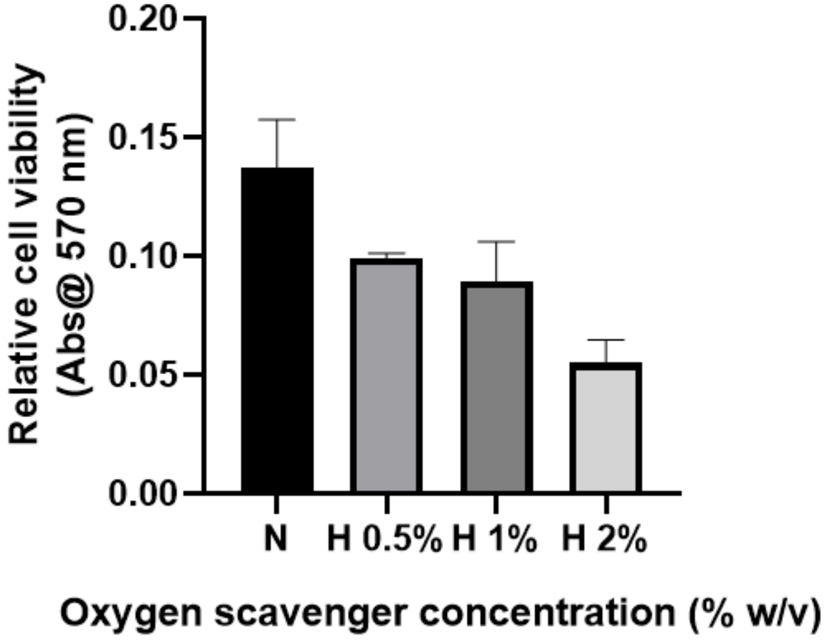


**Fig. S2.** Cell viability under different hypoxia conditions, assessed using Alamar Blue. Higher absorbance indicates greater cell viability and proliferation, while lower absorbance reflects increased cytotoxicity due to oxygen depletion by sodium sulfite over 24 hours

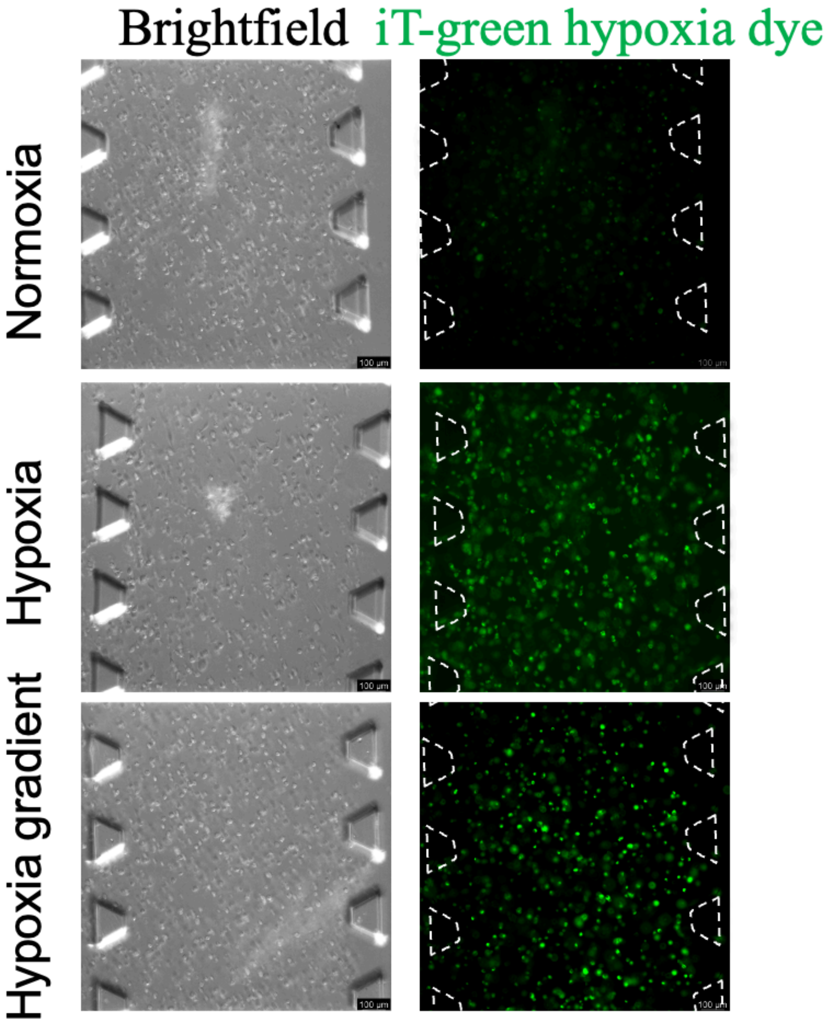


**Fig. S3.** Hypoxia monitoring for lung tumor on a chip model at three different hypoxia conditions, normoxia, hypoxia, and hypoxia gradient (second replicate).

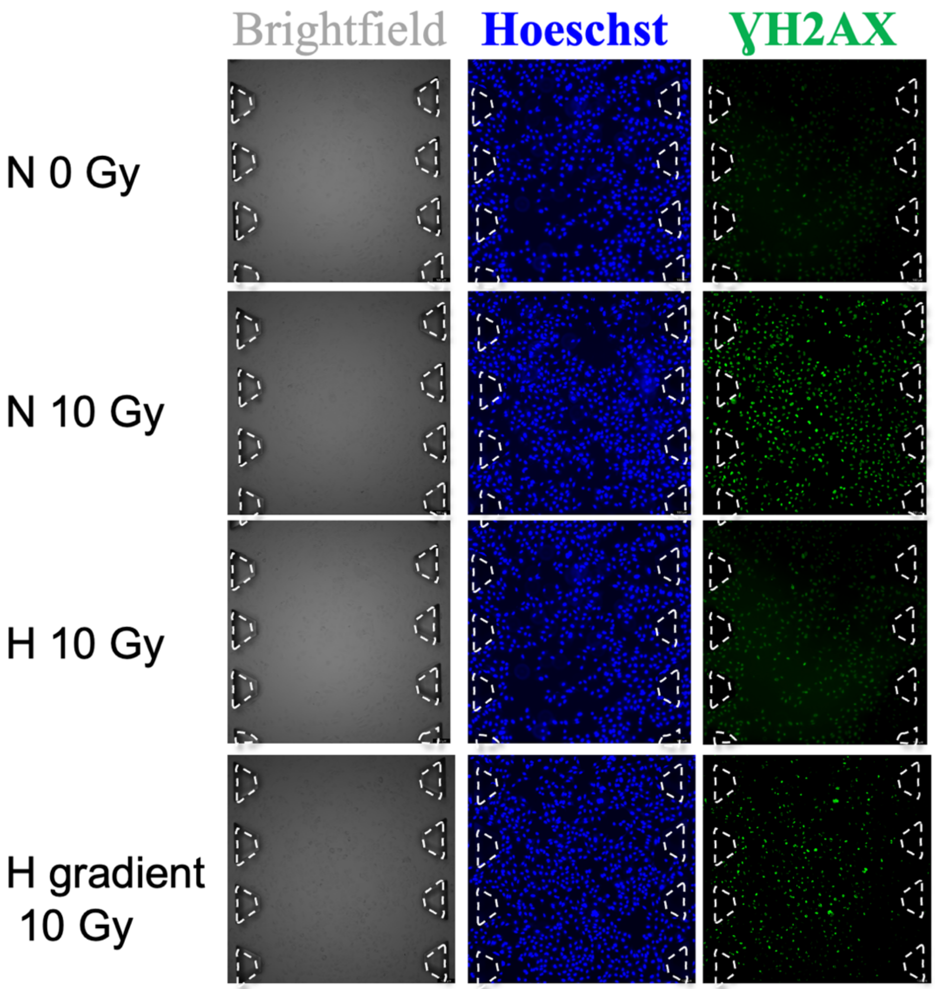


**Fig. S4.** DNA damage outcome for the lung tumor on a chip model using A549 cells (second replicate).

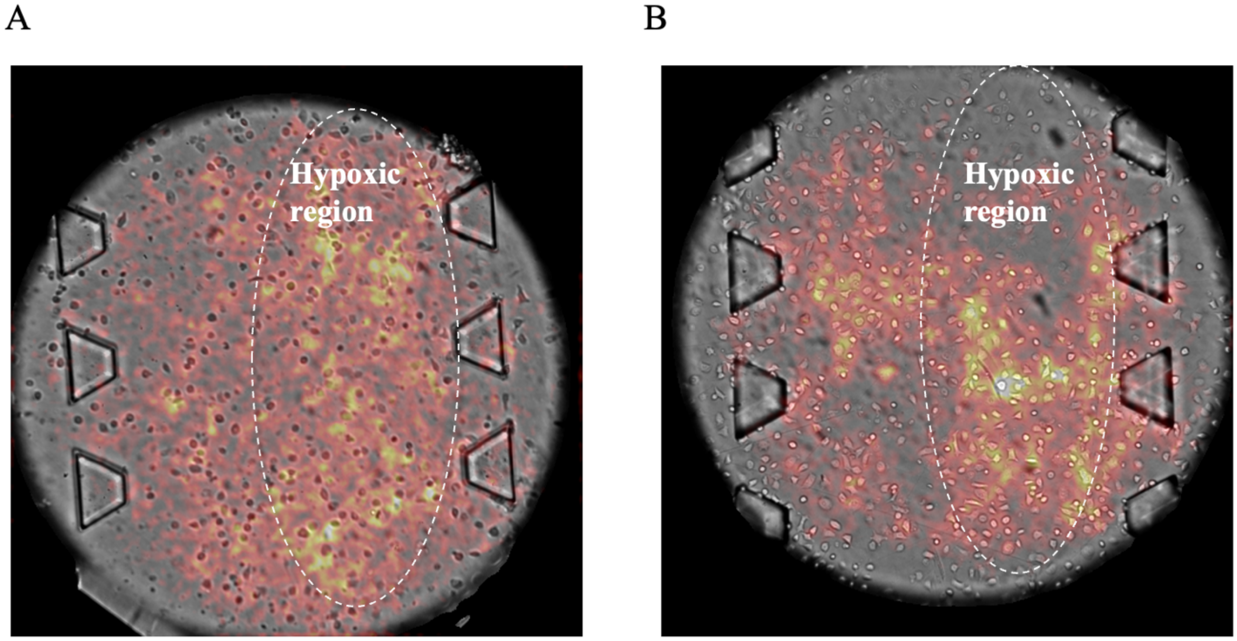


**Fig. S5.** RLM images showing FDG uptake in tumor-on-a-chip models containing A549 lung tumor cells grown in 2D (without hydrogel) under hypoxia gradient. Cells were cultivated directly on the surface of chips coated with attachment factor. (A) Chip 1, (B) Chip 2. Higher resolution can be achieved due to the closer proximity of the cells to the scintillator.

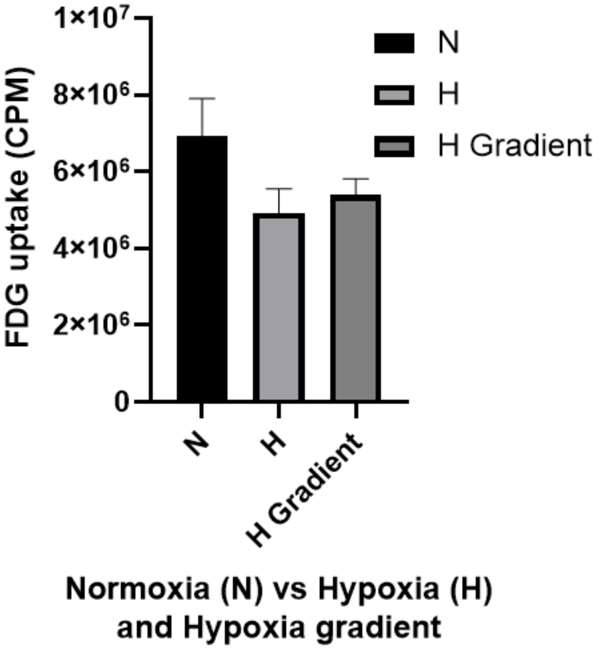


**Fig. S6.** FDG uptake by ToC models (based on A549 cells) before normalization by cell number.

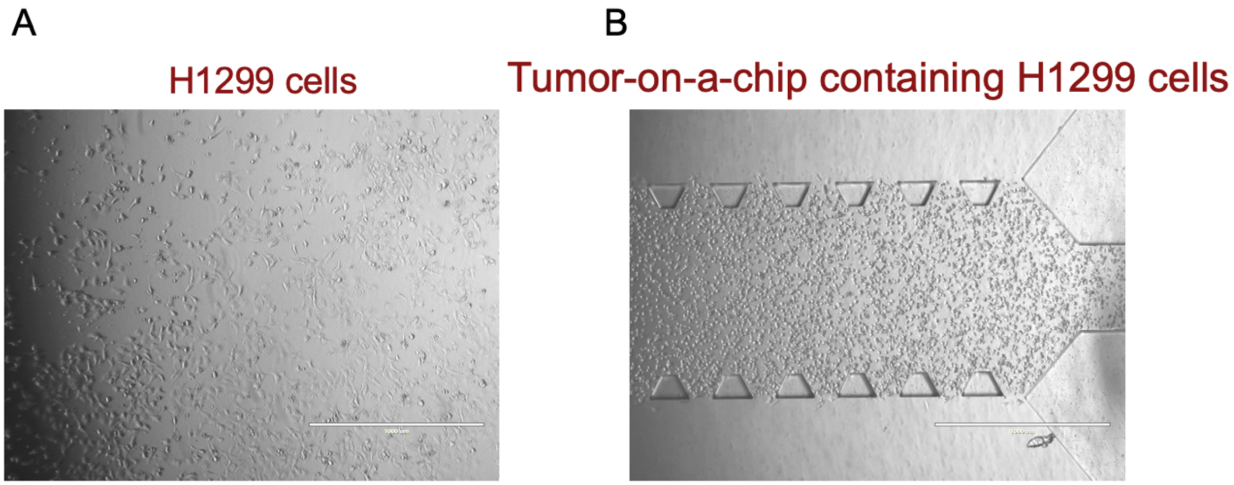


**Fig S7.** (A). H1229 cells in culture. (B). Tumor-on-a-chip containing H1299 cells grown in 3D in fibrin hydrogel (3 mg/ml).

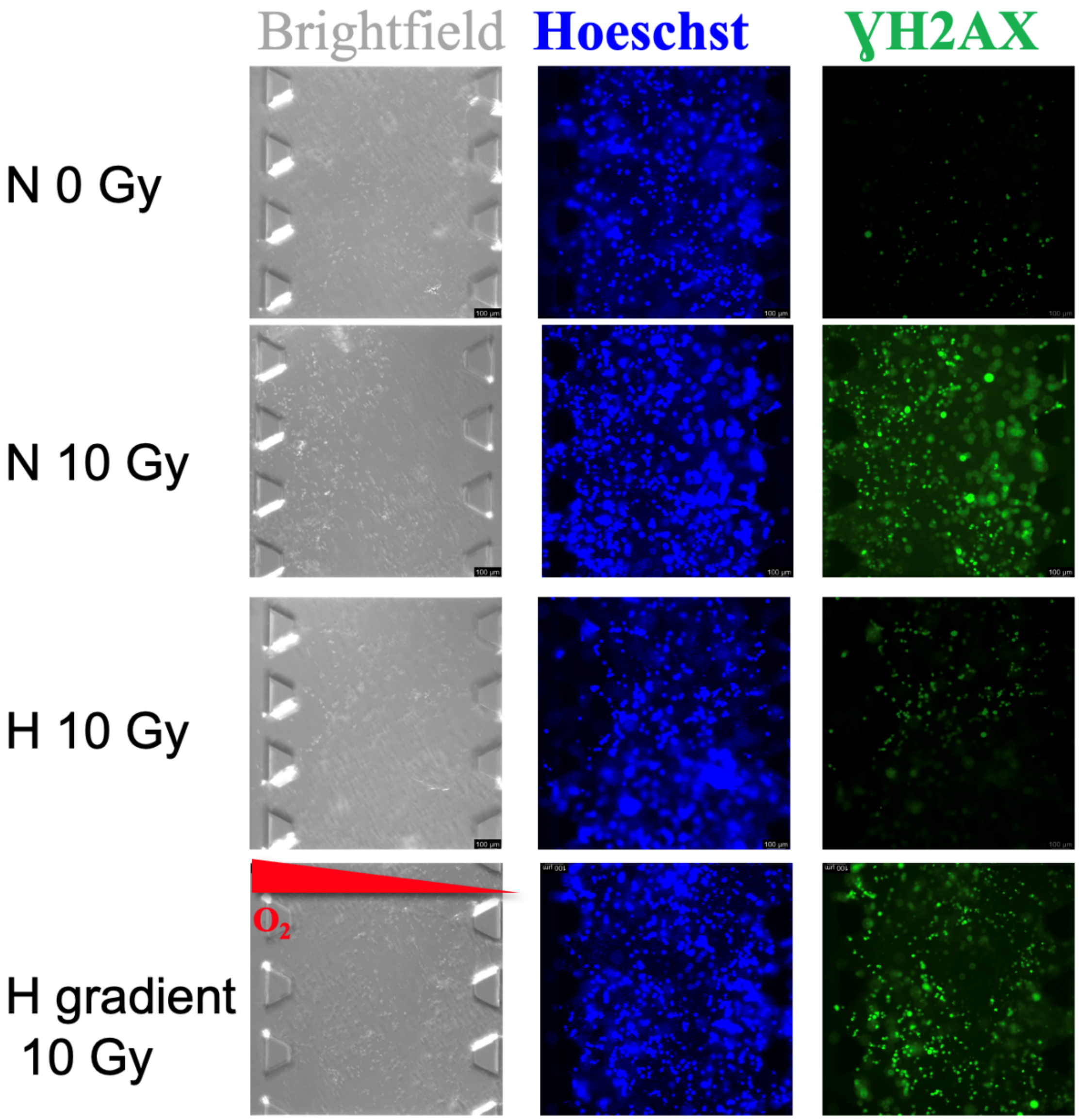


Fig. S8. Representative brightfield and fluorescence images (Hoechst and γH2AX staining) of H1299 tumor-on-a-chip models cultured under normoxia, hypoxia, and hypoxia gradient conditions after exposure to 10 Gy X-ray

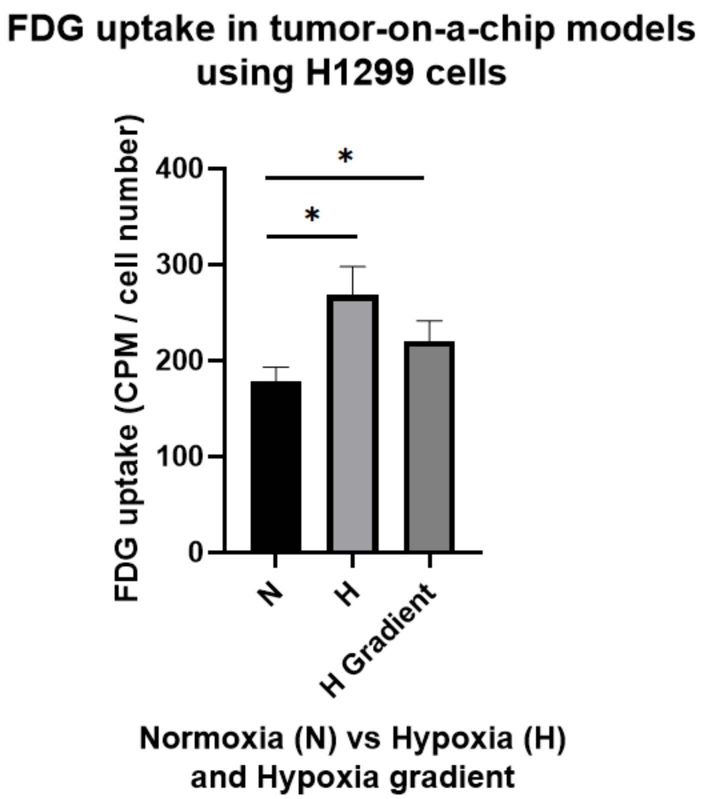


**Fig. S9.** FDG uptake by lung tumor-on-a-chip based on H1299 cells grown under normoxia, hypoxia, and hypoxia gradient. Data represent mean ± SD from $n = 3$ independent samples.
